# C25-modified rifamycin derivatives with improved activity against *Mycobacterium abscessus*

**DOI:** 10.1101/2021.07.12.452042

**Authors:** Laura Paulowski, Katherine S. H. Beckham, Matt D. Johansen, Laura Berneking, Nhi Van, Yonatan Degefu, Sonja Staack, Flor Vasquez Sotomayor, Lucia Asar, Holger Rohde, Bree B. Aldridge, Martin Aepfelbacher, Annabel Parret, Matthias Wilmanns, Laurent Kremer, Keith Combrink, Florian P. Maurer

## Abstract

Infections caused by *Mycobacterium abscessus* are difficult to treat due to its intrinsic resistance to most antibiotics. Formation of biofilms and the capacity of *M. abscessus* to survive inside host phagocytes further complicate eradication. Herein, we explored whether addition of a carbamate-linked group at the C25 position of rifamycin SV blocks enzymatic inactivation by Arr_Mab_, an ADP-ribosyltransferase conferring resistance to rifampicin. Unlike rifampicin, 5j, a benzyl piperidine rifamycin derivative with a morpholino substituted C3 position, is not modified by purified Arr_Mab_. Additionally, we show that the Arr_Mab_ D82 residue is essential for catalytic activity. Thermal profiling of Arr_Mab_ in the presence of 5j, rifampicin or rifabutin shows that 5j does not bind to Arr_Mab_. We found that the activity of 5j is comparable to amikacin against *M. abscessus* planktonic cultures and pellicles. Critically, 5j also exerts potent antimicrobial activity against *M. abscessus* in human macrophages and shows synergistic activity with amikacin and azithromycin.

## Introduction

*Mycobacterium abscessus* is a rapidly growing non-tuberculous mycobacterium (RGM) ^1^. In humans, Mab can cause severe pulmonary infections, particularly in patients with predisposing conditions such as bronchiectasis and cystic fibrosis ^2,3^. In addition, Mab can cause soft-tissue infections following surgery due to traumatic injuries or cosmetic procedures ^4^. Mab colonies show two phenotypically distinct morphotypes based on the presence or absence of glycopeptidolipids (GPL) in the mycobacterial cell wall. Smooth (S) variants express comparably high levels of GPL, whereas GPL production in rough (R) variants is significantly reduced or completely absent, due to irreversible mutations in genes involved in GPL biosynthesis or transport ^5,6^. GPL status and colony morphology play an important role in the interaction of Mab with the host and the environment by regulating biofilm formation and sliding motility, host–pathogen interactions and intracellular survival strategies, which dictate progression from colonization to disease and, ultimately, clinical outcomes ^3,5^. Mab is intrinsically resistant to most clinically available antibiotics and the success of current treatments for Mab pulmonary disease is below 50% ^2,7^. Standard multidrug treatment lasts for several months and is associated with a high risk of severe side effects including gastrointestinal distress, irreversible ototoxicity, and myelosuppression ^2,8^. Mab comprises three subspecies, *M. abscessus* subsp. *abscessus* (Mab_A_), *M. abscessus* subsp. *bolletii* (Mab_B_), and *M. abscessus* subsp. *massiliense* (Mab_M_) ^9^. Mab_A_ and Mab_B_ differ from Mab_M_ in that most clinical isolates are resistant to macrolides due to the presence of Erm(41), an inducible rRNA methylase ^10-12^. Macrolide resistance is a risk factor for worse clinical outcome ^13-16^. With increasing numbers of reported NTM infections and a global spread of highly virulent clones ^17^, new therapeutic options for Mab are urgently needed ^18^.

Rifampicin (RMP, Figure 1), which belongs to the rifamycin class of antibiotics, is a key drug for the treatment of tuberculosis ^19-22^. RMP targets the beta-subunit of the bacterial RNA polymerase (RpoB) ^23-25^ and exerts bactericidal activity ^26-28^ also against intra-macrophage ^29^ and non-replicating *M. tuberculosis* bacilli ^30^. Furthermore, rifamycins are known to diffuse within granulomas and biofilms, two niches that are also exploited by NTM ^31,32^. In Mab, resistance to rifamycins is conferred by a combination of several mechanisms including the intrinsically low permeability of the Mab outer membrane and presumably drug efflux pumps^33^. Previously, drug modification by an ADP-ribosyltransferase termed Arr has been demonstrated to play a crucial role in rifamycin resistance in *Mycobacterium smegmatis* (Msm; Arr_Msm_) ^34^. Arr_Msm_ acts by transferring an ADP-ribose unit from the donor (NAD^+^) to a susceptible amino acid residue on the target molecule with loss of nicotinamide ^35^. Recently, an Arr orthologue, Arr_Mab_ (encoded by MAB_0591), has been demonstrated to also confer resistance to RMP in Mab ^36,37^. In a Δ*arr*_Mab_ isogenic mutant of the Mab ATCC19977 type strain, the minimal inhibitory concentration of RMP dropped by 512-fold from 128 to 0.25 µg/mL as compared to the parental wild type strain, highlighting the importance of ADP-ribosylation for intrinsic resistance to RMP in Mab. Further research has shown that rifabutin (RBT, Figure 1), another rifamycin antibiotic, inhibits growth of all three Mab subspecies at concentrations of 3 to 9 µM (corresponding to 2.5 to 7.6 µg/mL) in Muller-Hinton medium ^38^. In addition, RBT shows bactericidal activity both on extracellular and intracellular forms of Mab and increases survival of Mab-infected zebrafish ^39^. However, using the Δ*arr*_Mab_ Mab ATCC19977 mutant, SCHÄFLE *et al*. recently demonstrated that despite its lower MICs compared to RMP, RBT remains partially modified by Arr_Mab 37_. These findings suggest that rifamycins can be further optimized to counteract ADP-ribosylation. Consequently, biochemical characterization of the interaction between Arr_Mab_ and different rifamycins is needed to allow for targeted drug design.

**Figure 1:**
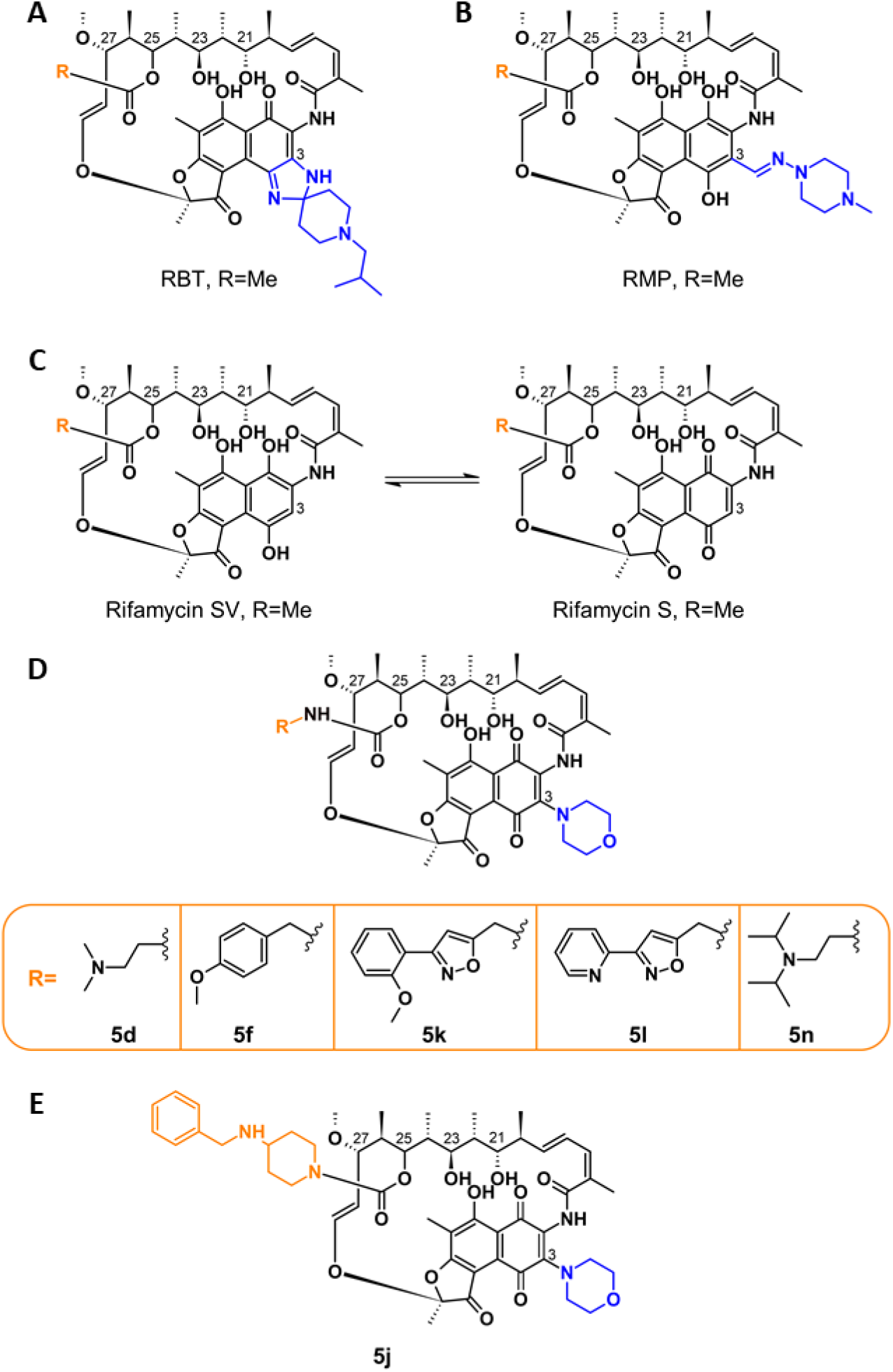
Structures of rifabutin (A), rifampicin (B), rifamycin SV and rifamycin S (C), and C25-modified rifamycin S derivatives used in this work (D). (Suppl. Table 1) ^34,40,41^.

Early work by COMBRINK *et al*. showed that C25 carbamate rifamycin derivatives are resilient to inactivation by ADP-ribosyl transferases in Msm ^34^. The authors found that relatively large groups attached to the rifamycin core *via* a C25 carbamate linkage prevented inactivation through ADP-ribosylation of the C23 alcohol catalyzed by Arr_Msm_. ROMINSKI *et al*. evaluated three of these compounds (5f, 5k, 5l; Figure 1D) against the Mab ATCC 19977 type strain and found modest activity with minimal inhibitory concentrations (MICs) of 2 to 8 µg/mL ^36^. Interestingly, C25 modification not only increased activity against the Mab type strain, but also against the Δarr_Mab_ mutant, indicating that the increased activity of C25 rifamycin derivatives is only partially due to resistance to modification by Arr_Mab_. Lastly, the authors concluded that the studied rifamycin derivatives were still partially inactivated by Arr_Mab_. In contrast, we recently found that compound 5j (Combrink et al. 2007, corresponding to 2g in Combrink et al. 2019, Suppl. Table 1), a benzyl piperidine rifamycin derivative with a morpholino substituted C3 position (Figure 1), showed lower average MICs (<0.5 µg/mL) than all previously investigated C25-modified rifamycin derivatives against clinical isolates of Mab_A_, Mab_B_, Mab_M 40_.

In this study, we report an in-depth characterization of the activity of 5j in comparison to other C25 rifamycin derivatives as well as to RMP and RBT. We aimed to explore whether rifamycin activity on Mab residing within host macrophages or within pellicles as a model for biofilm formation can be restored by addition of a carbamate linked group of sufficient size at the C25 position. We also addressed whether the effectiveness of ADP-ribosylation is dependent on species (Arr_Mab_, Arr_Msm_) or substrate (RMP, RBT, and 5j,).

## Results

### Compound synthesis

We synthesized six C25-modified derivatives based on the lead structure of rifamycin SV (Figure 1, Suppl. Table 1). Synthesis pathways for compounds 5d, 5f, 5j, 5k, and 5l were previously reported by COMBRINK *et al*. in US patent 7,250,413 B2 ^41^. In addition, compound 5n, which was first reported as compound 2f by COMBRINK *et al*. in 2019 ^40^, was added to the panel. The compounds differ in their degree of complexity and the number of hydrogen bond donors and acceptors (Figure 1). Compound 5j possesses a 4-(benzylamino)-1-piperidine group that is attached *via* a carbamate linker to position C25 of the core structure (Figure 1D). The C25 side group of 5j differs from the side groups of the other derivatives in that it exhibits one H-bond acceptor and no H-bond donor.

### 5j is active against M. abscessus grown in planktonic cultures and shows synergistic activity with amikacin and azithromycin

We tested the activity of the different rifamycin derivatives against the Mab ATCC 19977 reference strain and against a panel of clinical Mab, *M. chelonae* and *M. fortuitum* isolates representing the three RGM species most commonly associated with clinical disease (Figure 2). Minimum inhibitory concentration (MIC) values for the Mab reference strain (indicated by a red star in Figure 2) were found to be in the order of 2 to 32 µg/mL following three days of incubation, with the most potent compounds (including 5j) showing activity comparable to amikacin (AMK, MIC = 2 µg/mL). In the clinical isolates, 5j showed the highest activity followed by 5n, corroborating previous findings ^40^. As expected, MIC values for RMP were significantly higher (p < 0.0001). These results indicate that 5j has improved antimicrobial activity against clinical Mab isolates in comparison with other rifamycins, including RBT and RMP.

**Figure 2:**
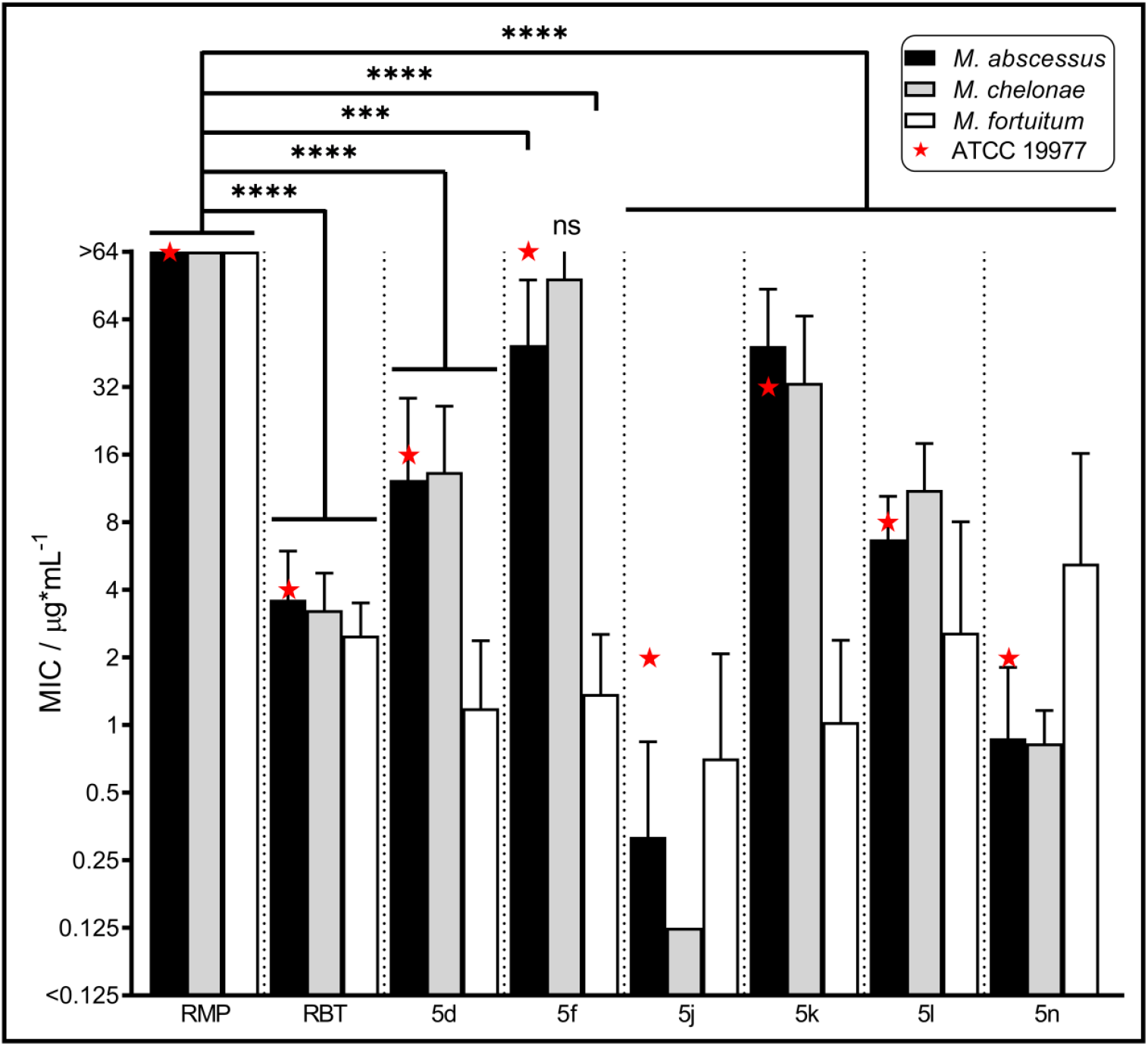
Minimal inhibitory concentrations of rifampicin, rifabutin, and rifamycin derivatives against *M. abscessus* ATCC 19977 and clinical NTM isolates. MICs were generated in cation-adjusted Muller-Hinton medium based on the procedure outlined in CLSI document M24 ^42^. MIC plates were read after 3 days of incubation at 30 °C. MIC values were obtained for *M. abscessus* (Mab_A_, n = 9; Mab_B_, n = 7; Mab_M_, n = 8), *M. chelonae* (n = 8), and *M. fortuitum* (n = 8). Data are plotted as mean ±SD. Data for the *M. abscessus* subsp. *abscessus* ATCC 19977 reference strain are depicted by (★). ***, P < 0.001; **** P < 0.0001; ns, not significant, confidence interval 99%.

We also explored whether 5j showed synergistic activity with other antimicrobials by applying the DiaMOND methodology that has been previously developed for *M. tuberculosis* ^43^. With log_2_FIC_50_ values of −0.75 for amikacin and −0.83 for azithromycin, 5j showed strong synergistic activity with both drugs against the Mab ATCC 19977 type strain. In contrast, no synergistic activity was observed for 5j with both linezolid (log_2_FIC_50_ = 0.61) and moxifloxacin (log_2_FIC_50_ = 0.38).

### 5j disrupts pellicle formation and decreases the number of viable cells in M. abscessus pellicles

Biofilm formation is thought to be one of the main features facilitating colonization and long-term survival of NTM within the host ^44^. We thus investigated if 5j or RBT showed activity against Mab ATCC 19977 in a pellicle formation assay adapted from OJHA *et al*. ^45^. After 72 h of incubation, a qualitative reduction in pellicle formation was observed following treatment with 5j, RBT, or AMK (Figure 3A). In contrast, pellicles treated with RMP retained a reticular formation at the air-liquid interface similar to the DMSO control (Figure 3A, red arrows). Corresponding CFU counts showed a 2- to 3-log reduction of viable cells upon treatment with 5j, RBT, or AMK whereas no effect was observed upon treatment with RMP or DMSO (Figure 3B). An example of a CFU count is shown in Figure 3D.

**Figure 3:**
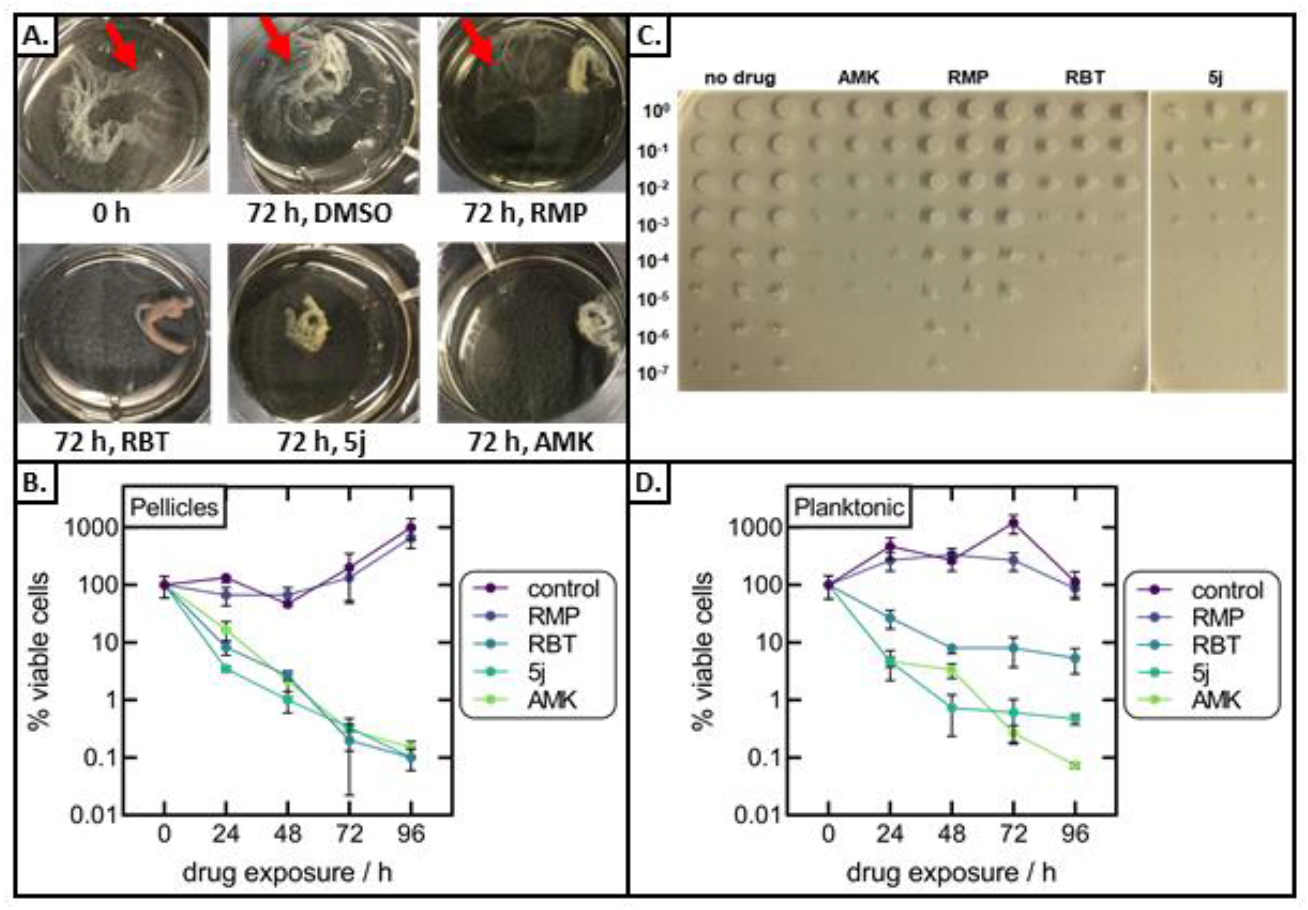
Inhibitory effect of rifamycins on *M. abscessus* pellicles and planktonic cultures. **A:** Appearance of Mab pellicles prior to drug exposure and following 72 h of exposure to 10 µg/mL of 5j, RBT, RMP or AMK. Pellicles treated with RMP retained a reticular formation at the air-liquid interface similar to the DMSO control (red arrows). **B:** Viable cell counts of Mab pellicles exposed to 5j, RBT, RMP, or AMK for up to 96 h without shaking. Data are plotted as mean ±SD of three independent experiments. CFU counts were normalized to the CFU count prior to the addition of antibiotics. **C:** Example of CFU counts from Mab pellicles obtained following 72 h of drug exposure. **D:** Viable cell counts in planktonic cultures incubated with shaking. Three technical replicates are shown per drug.

### 5j and rifabutin are active against M. abscessus within host phagocytes

Survival within host phagocytes is another hallmark of mycobacterial infections ^46,47^. We thus investigated if 5j or RBT exert intracellular antimicrobial activity against Mab smooth (S) and rough (R) morphotypes within THP-1 macrophages. Following infection at a multiplicity of infection (MOI) of 2:1 and killing of extracellular bacteria using 250 µg/mL AMK, macrophages were treated with either 5j (8 µg/mL), RBT (16 µg/mL), RMP (50 µg/mL) or AMK (50 µg/mL). DMSO treated macrophages were included as a negative control for intracellular bacterial replication. At 0, 1, and 3 days post-infection (dpi), macrophages were lysed and plated to determine intracellular bacterial loads. At 1 dpi, the presence of DMSO, RMP or AMK failed to reduce the intracellular burden of Mab S and R variants (Figure 4A and 4B). In contrast, exposure to 5j and RBT strongly decreased intracellular bacterial loads. At 3 dpi, further intracellular growth was observed as compared to 0 dpi for the DMSO and RMP treatments, while CFU counts remained stationary as compared to 1 dpi following treatment with AMK. In contrast, treatment with RBT or 5j led to a sustained reduction of intracellular bacteria for both S and R morphotypes indicating that both compounds kill both Mab variants within THP-1 cells. Of note, reduction of intracellular CFU was comparable for 5j and RBT although the concentration at which 5j was used was half the concentration of RBT (8 µg/mL and 16 µg/mL, respectively).

**Figure 4:**
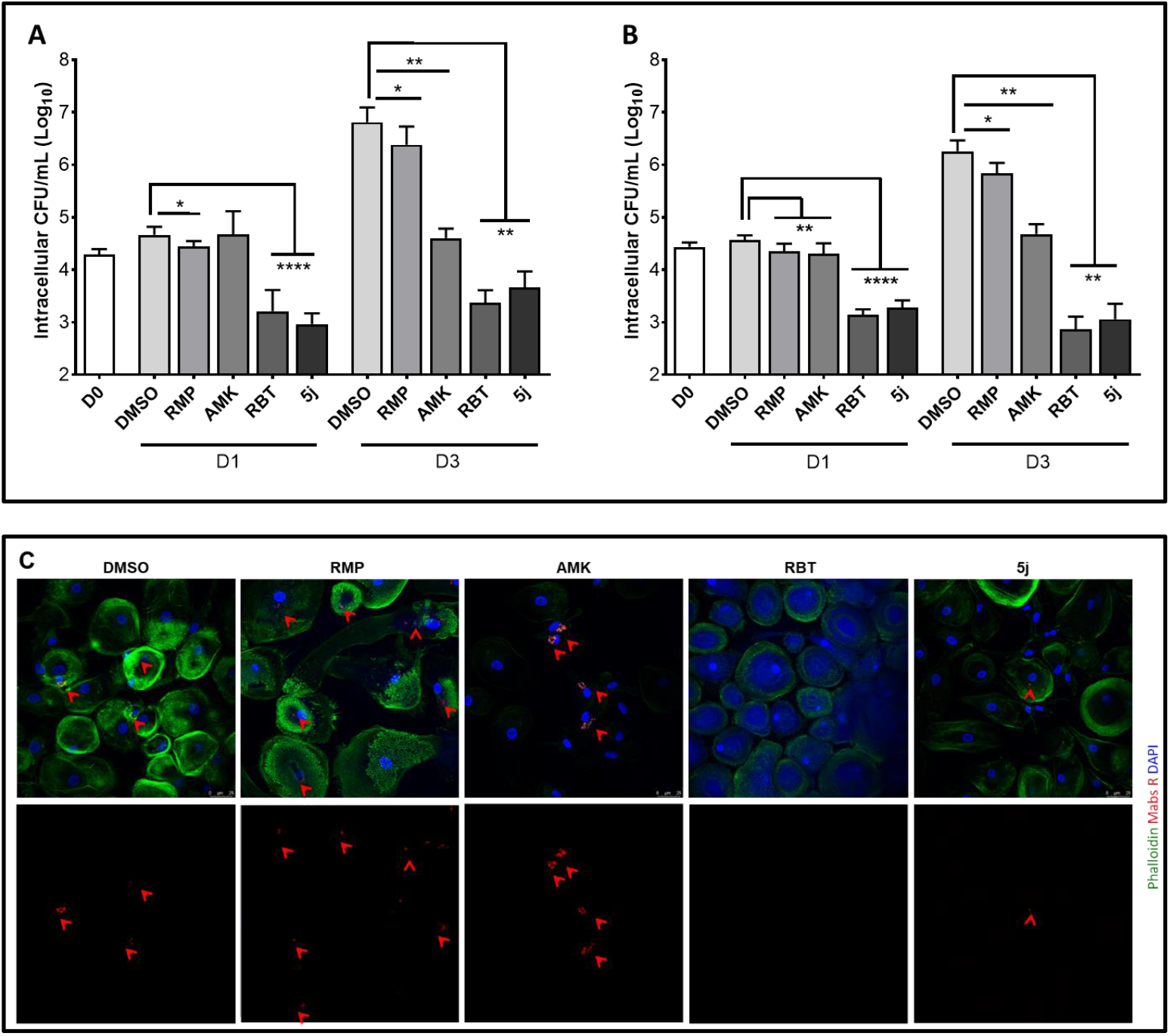
Intracellular survival of *M. abscessus* smooth and rough morphotypes within human macrophages treated with 5j, rifampicin, rifabutin or amikacin. THP-1 macrophages were infected with Mab CIP104536 smooth (**A**) or rough variant (**B**) at a MOI of 2:1 for 4 h. Data are plotted as mean ±SD of four independent experiments (n = 4). **C:** Confocal microscopy of human monocyte-derived macrophages (HMDM) infected for 6 h with Mab R expressing tdTomato at a MOI of 10:1. Bacterial cells appear in red and are additionally highlighted by red arrow heads. HMDM nuclei were stained with DAPI (blue). Actin staining was done with Phalloidin ^49^.

To further validate our findings, we performed fluorescence microscopy on primary human monocyte-derived macrophages (HMDM) infected with a Mab CIP 104536 R variant expressing tdTomato ^48^. HMDMs were grown in antibiotic-free cell culture medium and infected at a MOI of 10:1 for 6 h. After killing of extracellular bacteria with AMK, macrophages were treated with either 5j, RBT, RMP, AMK at a final concentration of 5 µg/mL each. Despite the higher MOI and lower drug concentrations, a pronounced reduction of red fluorescent bacilli was observed upon treatment with 5j or RBT at 3 dpi as compared to AMK or RMP treatment and the DMSO control, corroborating the quantitative results obtained in THP-1 cells (Figure 4C).

### Arr_Mab_ has no ribosylation activity on 5j

To investigate the enzymatic activity of Arr_Mab_ and Arr_Msm_ on different substrates, we purified both wild-type enzymes and an Arr_Mab_ mutant, in which we introduced the mutation D82A to the putative active site aiming to abolish enzymatic activity (Figure 5A+B) ^35^. The purified enzymes were incubated with NAD^+^, and 5j, RBT or RMP at room temperature. Reaction products were analysed by reversed phase thin layer chromatography (rpTLC; Figure 5C). When RMP was used as a substrate for Arr_Mab_ or Arr_Msm_, we observed a single orange band (reflecting the natural colour of RMP) in aliquots taken at t_0_. Then, after 5 min of incubation, reactions were resolved into two distinct bands. In parallel, the single band observed at t_0_ remained unchanged over time using ArrD82A_Mab_ and in control reactions without Arr or NAD^+^(Figure 5C). With RBT used as a substrate, a second spot also became discernible from 5 min onwards. However, the spot observed at t_0_ and in control reactions without Arr or NAD^+^ remained clearly present. When 5j was used as a substrate, no additional spots became visible throughout. Taken together, these results suggest that RMP was completely turned over by both Arr_Mab_ and Arr_Msm_ within 60 min, while RBT was partially modified and 5j was not modified.

**Figure 5:**
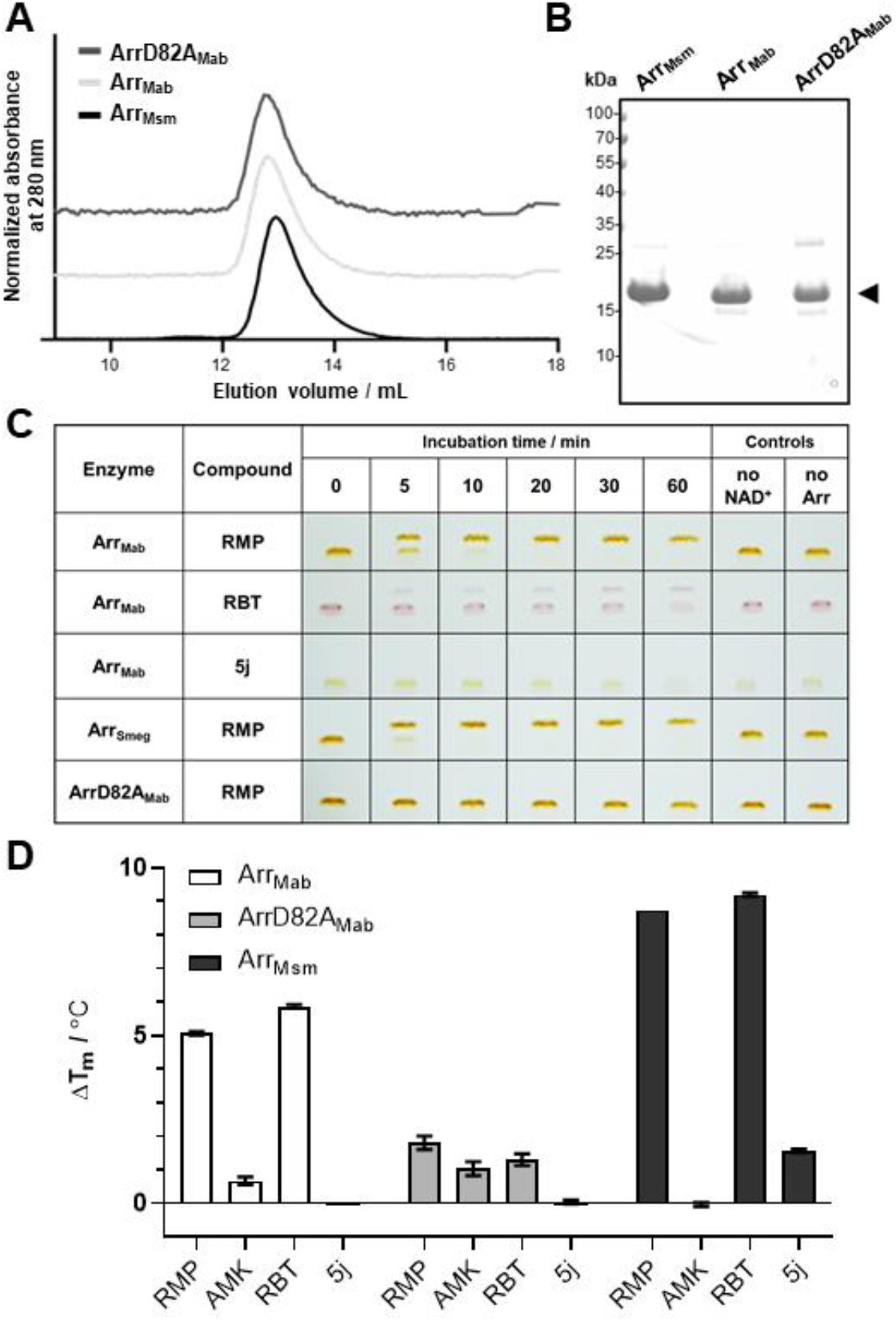
Interaction of purified Arr_Mab_, ArrD82A_Mab_, and Arr_Msm_ with rifampicin, rifabutin and 5j. **A:** Size exclusion profile of purified Arr_Mab_, the catalytically inactive mutant (ArrD82A_Mab_) and Arr_Msm_, with the corresponding peak fraction separated by SDS-PAGE. **B:** The arrow indicates the predominant band corresponding to the Arr variants. **C:** ADP-ribosylation reactions stopped after the indicated incubation times, analyzed by rpTLC. Reactions without NAD^+^ or without the enzymes were added as negative controls. **D**. Change in melting temperature (ΔT_m_) of Arr variants in the presence of selected compounds. The ΔT_m_ value was derived by subtracting mean T_m_ of the control (1% DMSO) from the mean T_m_ in the presence of ligand observed in NanoDSF. Melting experiments were performed in triplicate, RMP, 300 µM rifampicin; RBT, 300 µM rifabutin; AMK, 300 µM amikacin; 5j, 300 µM compound 5j.

To further investigate the binding affinity of 5j, RBT, RMP, and AMK as a negative control to Arr_Mab_, we subjected the purified Arr proteins to a thermal denaturation assay using Differential Scanning Fluorimetry (nanoDSF). In this approach, an increase in thermal stability of a protein indicates the stabilizing effect of a compound binding to it. Addition of either RMP or RBT led to an increase in the melting temperature (T_m_) of both Arr_Mab_ and Arr_Msm_ of at least 5 °C compared to the DMSO control (Figure 5D). This change indicates the binding of RMP and RBT to both Arr_Mab_ and Arr_Msm_. In contrast, no (Arr_Mab_) or only a slight (Arr_Msm_) increase in melting temperatures was observed when 5j was used as a substrate for both enzymes. In line with the rpTLC results, addition of none the drugs led to any increase in T_m_ with the active site ArrD82A_Mab_ variant, suggesting this mutation is very likely abolishing binding of rifamycin substrates to Arr_Mab_.

Notably, both rpTLC and nanoDSF experiments indicated a higher activity and stronger binding affinity of RMP with Arr_Msm_ as compared to Arr_Mab_. To investigate these differences, we performed isothermal titration calorimetry experiments to obtain insights into the thermodynamic binding parameters of Arr_Msm_. Titration of RMP into Arr_Msm_ or Arr_Mab_ revealed clear differences in the binding affinities of the two proteins. Arr_Msm_ displayed binding affinity to RMP in the nano molar range (290 ± 20 nM, Figure 6A), while Arr_Mab_ binding affinity to RMP was in the micro molar range (4.8 ± 0.003 mM, Figure 6B). To relate the binding affinity to the active site residues we generated a model of Arr_Mab_ using Arr_Msm_ in complex with RMP (PDB 2HW2) as a template ^35^. An overlay of the active sites revealed key differences in the residues involved in RMP binding (Figure 6C). In the Arr_Msm_ co-crystal structure Y37, F124 and F90 which are involved in hydrophobic interactions with RMP are replaced by F39, M126 and L92, respectively. This replacement results in an altered rifamycin binding pocket providing a structural rationale for the observed variation in ribosylation efficiencies.

**Figure 6:**
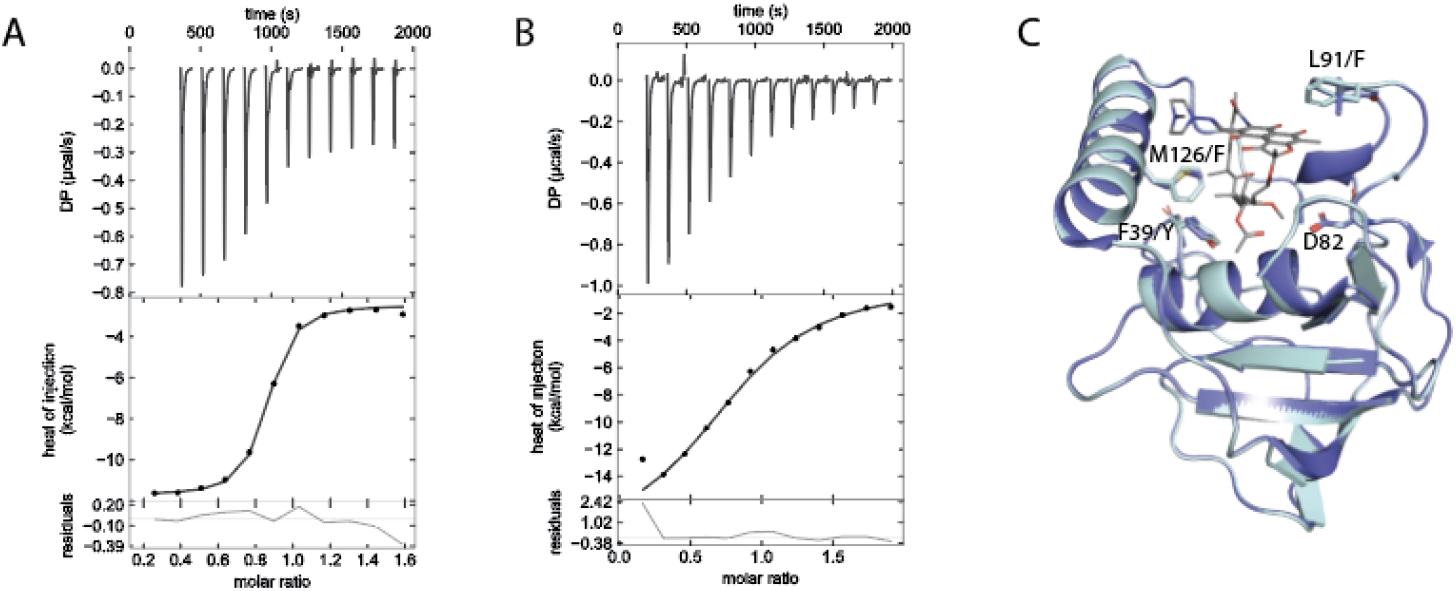
Arr active site and quantification of rifampicin binding. A, B: ITC thermogram of rifampicin titrated into Arr_Msm_ (A) and Arr_Mab_ (B). C: Overlay of Arr_Msm_ crystal structure (PDB 2HW2) with the Arr_Mab_ model, shown in dark blue and light blue, respectively. Residue numbering is according to Arr_Msm_. The catalytic aspartate (D82) has been shown in stick representation along with other residues which differ in the Arr_Mab_ RMP binding site. RMP is shown as a stick model.

## Discussion

The success of RMP in the treatment of tuberculosis has demonstrated that compounds inhibiting transcription can effectively disrupt proliferation of mycobacteria. Moreover, considering the activity of rifamycins on intracellular bacteria and biofilms, discovery or modification of rifamycins with activity against NTM has attracted renewed interest ^1,50,51^. Here, we showed that modification of the C25 position of rifamycin S by cross-linking *via* an imidazole carbamate active intermediate and the secondary amine 4-(benzylamino)-1-piperidine leads to improved antimicrobial activity of rifamycin S analogs against RGM, including clinical Mab isolates. The principal findings of our study are that 5j and RBT are significantly more effective than RMP against Mab *in vitro* and that this holds true not only for planktonic bacteria but also applies to bacteria contained within Mab pellicles or host phagocytes. Furthermore, our data regarding compound 5j demonstrate that ADP-ribosylation mediated by Arr can be further diminished as compared to both RMP and RBT by modification of the C25 position.

The range of MIC values reported in the literature for Mab and RBT shows considerable variation, which has been related to different growth media (CaMHB as recommended by CLSI versus Middlebrook 7H9 broth) and procedures, for example by adding alamarBlue to enhance readability of incubated BMD plates ^38,52^. In our hands, the antimicrobial activities of both 5j and RBT against Mab planktonic cultures were comparable to AMK, a cornerstone antibiotic in current multidrug regimens ^2^. Our findings also support the conclusion of AZIZ *et al*. that the MIC values of RBT, and as we show also those of 5j, are at least in the order of one log step lower than those of RMP ^38^. Of note, the RBT MIC values observed by AZIZ *et al*. for Mab_A_, Mab_B_, and Mab_M_ reference strains in CaMHB (3 to 9 µM, corresponding to 2.5 to 7.6 µg/mL) closely agree with our results for the Mab_A_ ATCC 19977 reference strain and the investigated Mab_A_, Mab_B_, and Mab_M_ clinical isolates (2 to 8 µg/mL). Strikingly, synergy studies revealed that 5j acts synergistically with both amikacin and also azithromycin even against Mab ATCC 19977 which shows inducible macrolide resistance due to a functional Erm(41) methyltransferase. In contrast, addition of 5j failed to potentiate the effect of linezolid and moxifloxacin, two orally administered drugs that are lacking sufficient activity against Mab alone.

A prevailing paradigm of chronic Mab infection is that biofilm formation contributes to persistence of Mab within the human host ^3^. In fact, Mab biofilm aggregates have been identified in the lungs of patients with CF, non-CF bronchiectasis, and COPD ^53,54^. In contrast, the Mab R variant is thought to represent an invasive, cord-forming phenotype emerging in the chronically infected host over time with the factors triggering S-to-R conversion by loss of surface GPL being incompletely understood ^3^. Biofilm aggregates are significantly more tolerant than planktonic variants to acidic pH, hydrogen peroxide and treatment with AMK or azithromycin ^55^. Our finding that RBT, and even more so 5j, penetrate into Mab pellicles indicate that these compounds could play a role in potentiating the activity of therapeutic regimens against bacteria contained in such aggregates both in early and late stages of the infection.

Likewise, activity of current standard regimens is insufficient against Mab bacteria contained within host phagocytes. GREENDYKE *et al*. found that both AMK and clarithromycin as well as cefoxitin, another commonly used first line drug to treat Mab infections, had only a bacteriostatic effect against bacteria contained within human macrophages ^49^. This agrees with early studies by HAND *et al*. demonstrating limited uptake of aminoglycosides and beta lactams into alveolar macrophages ^56^. Notably, while a pronounced intracellular accumulation of macrolides was demonstrated in the latter study, adaptive macrolide resistance of Mab due to the inducible Erm(41) presents an efficient mechanism to counteract this effect ^10^. For RMP, HAND *et al*. demonstrated an intracellular accumulation of 2- to 5-fold ^56^. Notably, RBT has an even higher intracellular penetration and tissue distribution as compared to RMP with *in vitro* intracellular/plasma concentration ratios in neutrophils and monocytes of 9:1 and 15:1, respectively ^57-60^. Consequently, our findings with RBT and 5j underline the potential usefulness of these compounds against intracellular bacteria, while RMP, which is efficiently inactivated by Arr, showed no such activity.

We have shown that ADP-ribosylation by Arr_Mab_ is different for RMP, RBT and 5j. Both TLC and thermal denaturation experiments confirmed lower affinity of Arr_Mab_ to RBT as compared to RMP. These assays also confirmed that addition of the bulky 4-(benzylamino)-1-piperidine at the C25 carbon atom of the rifamycin SV scaffold as in compound 5j further diminishes enzymatic modification by lowering the capacity of the rifamycin to stably bind to the enzyme, thereby avoiding modification. In addition to the C25 modification, the binding behavior of RMP, RBT and 5j to Arr is further influenced by the substituents at the C3 position. For RMP, a *N*-(4-methylpiperazin-1-yl)methanimine is attached to the C3 position of the ring system. This group behaves unlike the C3 side chain of RBT regarding its spatial and chemical properties. In the case of RBT, an 8-isobutyl-1,4,8-triazaspiro[4.5]dec-1-ene residue is located at C3. This residue with its spiro center adds an additional chiral center (axial chirality) to the core structure. The imidazolium of the triazaspiro residue donates electron density towards the spiro carbon and in turn the electron rich piperidine residue on the opposite yield a net electron density shift out of the ring system. This in turn influences the aromaticity and planarity of the π-system. With 5j, a conformationally similar morpholino-group was introduced at the C3 position harboring a more electronegative oxygen in its side chain as compared to RBT and RMP. Furthermore, 5j can undergo tautomerism at the core ring structure and is in redox equilibrium from hydroquinone to quinone structure (Figure 1C).

Based on the crystal structure of Arr_Msm_ and the importance of its D84 residue for catalytic activity, we hypothesized that the corresponding D82 residue of Arr_Mab_ would be similarly essential for the catalytic activity of the enzyme ^35^. Indeed, both TLC and thermal denaturation assays confirmed that mutation of D82 located within the substrate-binding loop of Arr_Mab_ (based on its structural similarity to Arr_Msm_) abolished ADP-ribosylation *in vitro*. This is compatible with a molecular mechanism where D82 stabilizes a developing oxocarbenium transition state enabling attack of the hydroxyl group at position C23 of the antibiotic to attack C1 of the ribose. D82 thus seems to play a role comparable to the invariant E in protein ADP-ribosyltransferases ^35^.

Open questions remain about bactericidal versus bacteriostatic activity of 5j and the threshold for acquired resistance to 5j and RBT in Mab, for example by genetic alterations in *rpoB*, which has been beyond the scope of this contribution. Furthermore, additional pharmacokinetics and pharmacodynamics studies with regard to the activity and toxicity of 5j in preclinical *in vivo* models, as well as on its oral bioavailability would be a natural next step. As part of such future studies, investigation of the degree of interaction of 5j with the human CYP3A4 and CYP2C9 cytochrome P450 enzymes, which are strongly induced by RMP, would be of particular interest. In conclusion, the data presented herein show that C25-modified rifamycins, such as 5j, deserve further study as they appear to be among promising new therapeutic options for infections caused by Mab.

## Materials & Methods

### Chemical synthesis

Synthesis of compounds 5d-f and 5j-n has been described previously ^41 34,40^ (Suppl. Table 1). In brief, commercially available rifamycin SV was used as starting compound and oxidized in a first step to rifamycin S. The hydroxyl groups at position C21 and C23 were protected by acetonide formation. Subsequently, a morpholino group could be introduced at the C3 position via nucleophilic aromatic substitution. The 3-morpholino-rifamycin S C21-C23 diol was protected as an acetonide followed by hydrolysis of the C25 acetate to produce the corresponding C25 alcohol. The C25 carbodiimide (CDI)-adduct and the introduction of the benzyl group were both prepared according to US patent US 7,250,413 B2 ^41^. For amine displacement, an excess of 4-(benzylamino)-1-piperidine was added. The crude product was oxidized with potassium hexacyanoferrate, washed once with sodium bicarbonate and several times with sodium chloride. After drying over sodium sulfate the product was isolated in the quinone form which could be further purified via flash chromatography in acetone/dichloromethane supplemented with 1-5% MeOH for polar compounds, respectively. Reverse phase C18 modified silica gel was used as the stationary phase. All chemicals and consumables were purchased through Acros Organics (ThermoFisher, Scientific, Rochester, NY, USA) and Merck (Merck KgaA, Darmstadt, Germany).

### Antimicrobial susceptibility testing

MICs were determined by the broth microdilution method adapted from Clinical and Laboratory Standards Institute document M24 ^42^. In brief, an inoculum suspension was prepared in demineralized water by collecting growth from pure bacterial cultures using sterile cotton swabs and photometrically adjusting the optical density to a McFarland 0.5 turbidity standard. Following further dilution, the inoculum (c_final_ = 1-5⨯ 10^5^ CFU/mL) was then added to sterile 96-well plates (Greiner, Frickenhausen, Germany) containing serial antibiotic dilutions of either synthesized compounds or RMP, RBT, or AMK (Sigma-Aldrich, St. Louis, MI, USA) dissolved in DMSO (c_final_ <1%) and diluted in Cation-adjusted Muller-Hinton broth (CaMHB, Sigma Aldrich) without OADC. Plates were covered with adhesive seals, incubated at 30 °C in ambient air and examined for growth after 3, 5 and 7 days of incubation. For each isolate, an appropriate dilution of the inoculum suspension was plated on boiled blood and 7H10 agar plates (Becton Dickinson, Heidelberg, Germany) to control for purity of the bacterial culture and correct inoculum density.

### Drug interaction measurement

Diagonal measurement of n-way drug interactions (DiaMOND) was used to measure drug interactions ^43,61^. Drug powders were obtained from Sigma and dissolved in DMSO. Single-use aliquots were used to perform the assay. *Mycobacterium abscessus* strain ATCC 19977 was cultured in 7H9 medium supplemented with 0.05% Tween 80, 0.2% Glycerol, and 10% BBL Middlebrook ADC. Cultures were started from frozen aliquots and allowed to grow to mid-log phase with shaking at 37 °C overnight. Cultures were then sub-cultured before performing assays. Drugs were dispensed using an HP D300e digital dispenser. Bacteria were diluted to OD_600_ = 0.05, and 50 μL of diluted culture was added to clear 384-well plates. Plates were sealed with an optically clear plate seal and incubated in a 37 °C standing incubator for 48 h. OD_600_ was read using a standard plate reader. The concentration at 50% growth inhibition (IC_50_) obtained from the combination dose-response curves (observed IC_50_) and the IC_50_obtained from the estimation from the single drug dose-response curves (expected IC_50_) were used to calculate the fractional inhibitory concentration at the IC_50_ (FIC_50_) using Loewe Additivity ^43^. Log-transformed FIC_50_ values are reported such that log_2_FIC_50_ < 0 or > 0 indicate synergistic and antagonistic interactions, respectively. Drug interaction scores are reported as the mean of biological triplicate experiments.

### Pellicle formation assay

Antimicrobial susceptibility testing of Mab pellicles was adapted from OJHA *et al*. ^45^. Two Erlenmeyer flasks, each containing 20 mL CaMHB without OADC, were inoculated with one colony of Mab ATCC 19977 and incubated for 3-4 days at 37 °C with shaking (100 rpm). Then, sterile 12-well cell culture plates were filled with 1 mL CaMHB per well and inoculated with 10 µL of saturated planktonic Mab culture. Plates were incubated for 3 days at 37 °C with (planktonic culture for comparison) or without (pellicle formation) shaking at 100 rpm. Following incubation, RMP, RBT, 5j, or AMK were added at a final concentration of 10 µg/mL. Drug-free controls containing only DMSO were set up in parallel. The plate pairs were incubated for another 96 h. Images of the macroscopic pellicle appearance were taken at 72 h post-inoculation (hpi). At 0, 24, 48, 72, and 96 hpi, growth in one well incubated with and without shaking, respectively, was homogenized and transferred to 2 mL Eppendorf tubes. Cells were pelleted, washed two times with 500 µL CAMHB to remove residual antibiotics, and appropriate dilutions were plated for CFU counting on LB plates. All experiments were performed in triplicate.

### Intracellular activity in infected THP-1 monocytes

THP-1 macrophages were infected with S or R variants of Mab CIP104536 expressing Tdtomato at a MOI of 2:1 (bacteria:macrophage) for 4 h, after which macrophages were washed three times with PBS and treated with RPMI supplemented with amikacin 250 µg/mL for 2 h to kill extracellular bacteria. After treatment, macrophages were washed three times with PBS and then incubated with RPMI supplemented with either DMSO, rifampicin (50 µg/mL), amikacin (50 µg/mL), rifabutin (16 µg/mL) or 5j (8 µg/mL). Each day, macrophages were washed twice with PBS and drugs were replenished. When required, macrophages were lysed using 1% Triton X-100 in PBS, made up to a total of 1 mL with PBS and plated on LB agar to determine intracellular CFU.

### Immunofluorescence in human monocyte-derived macrophages

Human monocyte-derived macrophages (HMDM) were isolated from peripheral blood buffy coats as described previously ^62^. Cells were cultured in RPMI1640 containing 20% autologous serum and penicillin/streptomycin with weekly medium changes and used for infection after 1 week. The medium was replaced with cell culture medium without any antibiotics one day before infection. Construction of Mab mutants expressing tdTomato (λ_Ex,max_ = 554 nm; λ_Em,max_ = 581 nm) was reported previously ^48^. Bacteria were grown aerobically at 37 °C in Middlebrook 7H9 medium supplemented with 250 µg/mL Hygromycin B (Sigma-Aldrich) and harvested after 72 h by centrifugation followed by resuspension in ice-cold PBS containing 1 mM MgCl_2_ and CaCl_2_ (Sigma-Aldrich, St. Louis, MO, USA). Bacterial suspensions were homogenized by 50 passages through a syringe with a 25G needle and measured for optical density. Bacterial cells were then added to HMDMs at a multiplicity of infection (MOI) of 10:1 for 6 h. After infection cells were washed with PBS three times and incubated with RPMI medium containing 250 µg/mL AMK for 1 h, washed again with PBS and incubated with RPMI containing 50 mg/mL AMK for another 16 h to eliminate extracellular bacteria. Then cells were washed twice and incubated with RPMI containing 5j, RBT, RMP, AMK at a final concentration of 5 µg/mL or DMSO alone. After 72 h, cells were plated on coverslips (Marienfeld GmbH, Lauda-Königshafen, Germany), fixed with 3.7% (v/v) formaldehyde in PBS for 5 min, permeabilized with 0.1% Triton X-100 in PBS for 5 min, and incubated in blocking solution with 5% BSA in PBS for at least 15 min. Staining was performed with a 1:200 dilution of Alexa488-labeled phalloidin (Invitrogen, Carlsbad, USA) and a 1:500 dilution of 300 nM DAPI (Invitrogen, Carlsbad, USA) for 45 minutes. After three wash steps in PBS, coverslips were mounted in Mowiol (Calbiochem, Darmstadt, Germany) and analyzed by confocal microscopy. Images were acquired with a confocal laser scanning microscope (Leica DMI 6000 with a Leica TCS SP8 AOBS confocal point scanner) equipped with a 63x oil immersion HCX PL APO CS objective (NA 1.4–0.6). Acquisition was completed with Leica LAS AF software (Leica Microsystems, Wetzlar, Germany) and Volocity 6 software (PerkinElmer Life Sciences, Boston, USA) was used to process images.

### Cloning and expression of constructs

*MAB_0591* and *MSMEG_1221* were amplified by PCR from Mab ATCC 19977 or Msm mc^2^155, respectively, using Q5 High Fidelity Polymerase (New England Biolabs) using the primers listed in Suppl. Table 2. The expression vector pCoofy1 was linearized using NcoI/NotI to remove the *cddB* gene while maintaining the N-terminal His6 tag proceeded by a C3 precision protease sequence. DNA fragments were ligated into pCoofy1 using SliCE ^63^, generating pCoofy1-MabsArr and pCoofy1-MsmegArr. Site directed mutagenesis of pCoofy1-MabsArr by PCR using non-overlapping primers (Suppl. Table 2), generating the ArrMab inactive mutant (pCoofy1-MabsArrD82A). All expression constructs were sequence verified. The *E. coli* strain DH5a was used for all cloning experiments.

### Protein expression and purification

Expression plasmids were transformed into *E. coli* BL21(DE3). Cells were cultured in TB medium at 37 °C to an OD_600_ of 0.6 and protein expression was induced with 0.5 mM isopropyl-β-D-thiogalactopyranoside (IPTG). Cells were cultured for a further 18 h at 20 °C and pelleted by centrifugation, cell pellets were stored at –20 °C until required. Prior to cell lysis cells were resuspended in Lysis Buffer (300 mM NaCl, 20 mM HEPES, 20 mM imidazole supplemented with 1:100 protease inhibitor mix HP (SERVA Electrophoresis GmbH, Heidelberg, Germany) and 0.01% deoxy-ribonuclease I (Sigma-Aldrich, St. Louis, MO, USA) adjusted to pH 7.4. Cells were lysed using an Emulsiflex C3 high pressure homogenizer (AVESTIN, Ottawa, ON, Canada) by performing three cycles of 15,000 psi at 4 °C. The cell suspension was centrifuged (20 min, 43,000 ⨯ *g*, 4 °C) to pellet cell debris and passed through a 0.45 µm filter. Cell lysate was loaded onto a 5 mL HisTrap HP (GE Healthcare, Chicago, IL, USA) equilibrated with Buffer A (300 mM NaCl, 20 mM HEPES, 20 mM imidazole; pH 7.4). The loaded column was washed with 20 column volumes (CV) of Buffer A and eluted using a linear gradient of up to a final concentration of 500 mM imidazole. Samples were analyzed by SDS-PAGE and fractions containing Arr were pooled and C3 precision protease added at a molar ratio of 100:1 (Arr:C3). The Arr:C3 mix was dialyzed against SEC buffer (100 mM NaCl, 50 mM HEPES; pH 7.4) overnight at 4 °C. The cleaved Arr sample was loaded on to a 5 mL HisTrap HP and the flow through, containing the cleaved Arr protein was pooled and concentrated using 3000 Da MWCO concentrators (Amicon). Concentrated Arr samples were injected into a Superdex 75 16/60 size-exclusion chromatography (SEC) column (GE Healthcare, Chicago, IL, USA) pre-equilibrated in SEC buffer for removal of aggregated protein. Eluted fractions were analyzed by SDS-PAGE and compared to the PageRuler Prestained Protein Ladder (Thermo Fisher Scientific, Waltham, MA, USA) for size approximation.

### Reversed-phase thin layer chromatography

TLC experiments were adapted from ^64,65^. In brief, purified enzymes (2 µM) were incubated with NAD^+^ (10 mM) and 5j, RBT or RMP (1 mM) in dH_2_O containing 50 mM [Tris(hydroxymethyl)methylamino]propanesulfonic acid (TAPS), 10 mM MgCl_2_, 1 mg/mL bovine serum albumin, and 20 mM dithiothreitol (DTT) at room temperature. Aliquots were taken at 0, 5, 10, 20, 30, and 60 min, stopped with equal volumes of ice-cold MeOH containing 1 M CO(NH_2_)_2_, and spotted on partially octadecyl (C18) modified silica plates (Norcross, GA, USA). As liquid phase, CHCl_3_:MeOH was used at a ratio of 2:1 (v/v) to separate the reaction products on TLC membranes for 10 min without staining.

### Thermal denaturation assay

Purified Arr proteins were diluted with SEC buffer to a concentration of 30 μM for analysis by NanoDSF. The compounds RMP, RBT, AMK and 5j were added to a final concentration of 300 μM, equivalent to a final concentration of 1% DMSO which was tested as a control. Samples were loaded into standard grade NanoDSF capillaries (Nanotemper) and loaded into a Prometheus NT.48 device (Nanotemper) controlled by PR.ThermControl (version 2.1.2). An excitation power of 50% was used to obtain fluorescence readings above 2,000 RFU for F330 and F350. Samples were heated from 20 °C to 90 °C with a slope of 1 °C/min. Melting experiments were done in triplicate.

### Modelling of Arr_Mabs_

The homology model of Arr_Mabs_ was generated using SWISS-MODEL ^66^ using default parameters with Arr_Msm_ crystal structure in complex with RMP (2HW2) as a template.

### Isothermal Titration Calorimetry

Purified Arr_Msm_ and Arr_Mab_ were prepared as described above and were extensively dialyzed against SEC buffer (100 mM NaCl, 50 mM HEPES; pH 7.4) overnight at 4 °C. All ITC reactions were performed at 25 °C using the MicroCal PEAQ-ITC. 30 μM of either Arr_Msm_ or Arr_Mab_ was loaded into the cell with 250 μM or 300 μM RMP, respectively, placed into the syringe. Heats of binding for all the reactions were integrated using NITPIC ^67^, fitted using a single-site binding model with SEDPHAT ^68^ and plotted with GUSSI ^69^. All titrations were performed as triplicates and errors are reported as standard deviations (SD).

### Statistical analysis

Data were evaluated using a two-tailed student’s unpaired t-test (Figure 2, GraphPad Prism Version 9) with a confidence interval of 99% or using a student’s paired t-test (Figure 4, GraphPad Prism Version 6.0). Significance values are as follows: non-significant: ns P > 0.05; significant: * P ≤. 0.05; very significant: ** P ≤ 0.01; extremely significant: *** P ≤ 0.001; **** P ≤ 0.0001.

## Acknowledgments

The authors are grateful for the expert technical assistance of Anja Goedicke at University Medical Center Hamburg and the staff at the German National Reference Laboratory for Mycobacteria throughout this study. We acknowledge technical support by the SPC facility at EMBL Hamburg. We thank Frank Bentzien (University Medical Center Hamburg-Eppendorf) for provision of peripheral blood buffy coats, and Dr. Antonio Virgilio Failla and Bernd Zobiak (UKE Microscopy Imaging Facility, University Medical Center Hamburg-Eppendorf) for their excellent assistance in performing immunofluorescence experiments.

## Author contributions

KC and FPM conceived the study. KC synthesized the investigated rifamycin derivatives. KB, LA, MDJ, LB, FPM obtained experimental data. NV, YD, and BBA designed, performed, and analyzed the drug interaction studies. HR and MA provided expert advice on biofilm assays, monocyte infection and immunofluorescence imaging. KB, SS, AP and MW designed, cloned and purified Arr enzymes and obtained affinity measurements. LP, KB, MDJ, LB, FVS, AP, MW, LK, KC and FPM analyzed the data. LP, FVS and FPM drafted the initial version of the manuscript. All authors contributed to the final version of the manuscript and approved its submission.

## Supplementary material

**Supplementary Table 1:** Compound designations and reference for synthesis methodology.

| Compound | Designation in<br>Combrink et al. 2019 | Substituent R1 | Synthesis | NMR | MS |
| --- | --- | --- | --- | --- | --- |
| 5f | 2a | 1-methoxy-4-methylbenzene | 34,41 | [34] | [34] |
| 5g | 2b | *N,N*,4-trimethylaniline | 34,41 | [34] | [34] |
| 5j | 2g | 4-(benzylamino)-1-piperidine | 34,41 | [34] | [34] |
| 5k | 2d | 5-ethyl-3-(2-methoxy)phenylisoxazole | 34,41 | [34] | [34] |
| 5l | 2e | 2-(5-ethyl-3-isoxazolyl)pyridine | 34,41 | [34] | [34] |
| 5n | 2f | Propyldiisopropylamine | 34,41 | [34] | [34] |

**Suppl. Table 2:**
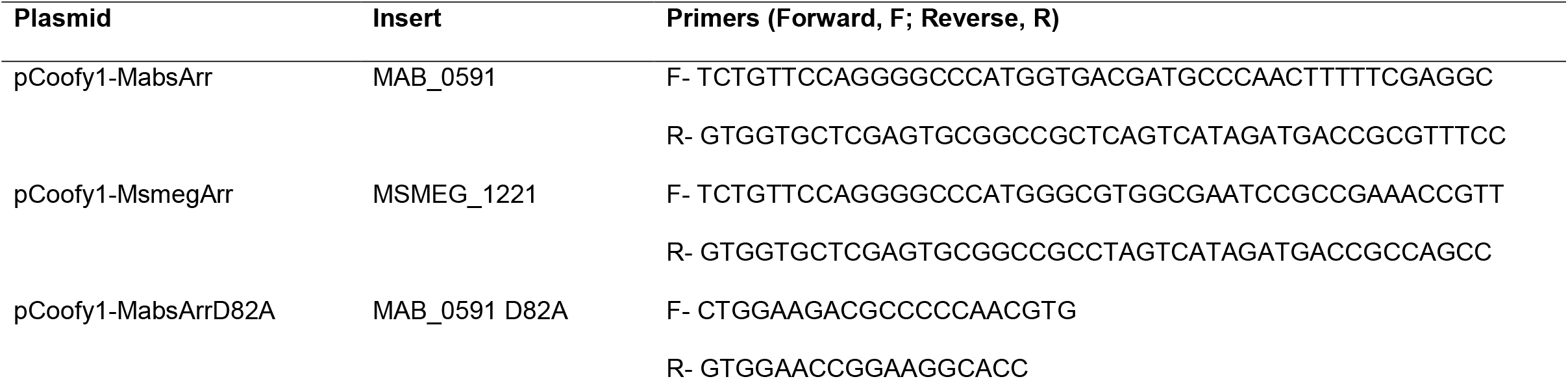
Plasmids and primers used to clone Arr_Mab_, Arr_Msm_, and ArrD82A_Mab_.

